# An Integrated Chemical Proteomics Approach for Quantitative Profiling of Intracellular ADP-Ribosylation

**DOI:** 10.1101/568527

**Authors:** Karunakaran Kalesh, Saulius Lukauskas, Aaron J. Borg, Ambrosius P. Snijders, Anthony K. L. Leung, Dorian O. Haskard, Peter A. DiMaggio

## Abstract

ADP-ribosylation is integral to a diverse range of cellular processes such as DNA repair, chromatin regulation and RNA processing. However, proteome-wide investigation of its cellular functions has been limited due to numerous technical challenges including the complexity of the poly(ADP-ribose) (PAR) chains, low abundance of the modification and lack of sensitive enrichment methods. To facilitate live-cell profiling of ADP-ribosylated proteins, we show that 2-alkyne-adenosine (2YnAd) is metabolically incorporated in mammalian cells and enables fluorescent detection and robust affinity enrichment of the modified proteins. We then present an integrated chemical biology approach that involves simultaneous metabolic incorporation of 2YnAd and the previously reported 6-alkyne-adenosine (6YnAd) in live cells followed by click chemistry with a capture reagent to facilitate highly sensitive and comprehensive enrichment of the modified proteins. By combining this dual metabolic labelling strategy with the tandem mass tag (TMT) isobaric mass spectrometry, we have quantified the responses of thousands of endogenous proteins to clinical PARP inhibitors Olaparib and Rucaparib. Our study provides insight into the wider scope of PARP targets and will help to advance further characterisation of their functional roles.

## Introduction

Protein ADP-ribosylation is a post-translational modification (PTM) where members of a family of enzymes known as ADP-ribosyltransferases (or PARPs) covalently link ADP-ribose moieties derived from NAD^+^ to the side chains of several possible amino acid residues (glutamate, aspartate, arginine, lysine, cysteine, serine) on target proteins. This PTM exists either as a monomer (mono-ADP-ribosylation), consisting of a single ADP-ribose moiety on the target residue, or as a polymer (poly-ADP-ribosylation) where a branched polymeric chain of ADP-ribose units originates from the target residue. Of the 17 PARPs encoded in the human genome, PARPs 1, 2 and 3 are known to have important roles in DNA regulation, particularly in the base excision repair pathway^1^. Consequently, PARP inhibitors have been developed as cancer therapeutics to exploit the concept of synthetic lethality in patients with germline mutations in *BRCA1* or *BRCA2* genes^2^. Beyond the established roles of these nuclear PARPs in DNA damage response, the broad cellular functions for the majority of the other members of the PARP family remain elusive primarily due to the lack of analytical techniques for the large-scale profiling of intracellular ADP-ribosylation.

Mass spectrometry (MS) proteomics studies of ADP-ribosylation in particular have been limited due to numerous technical challenges. These include low specificity and/or affinity when using recombinant macro domains^3,4^, *cis*-diol binding boronate beads^5^, antibodies^6^ or adapted phosphoproteome enrichment strategies^7^ to enrich for this low abundance PTM against complex cellular backgrounds. Additionally, most methods reported to date require *a priori* knockdown of poly-ADP-ribose glycohydrolase (PARG) activity to sufficiently increase the baseline level of poly-ADP-ribosylation for MS detection, which is not ideal since PARG knockdown is known to induce physiological changes in cells^8^. A further limitation of many previous studies is that they have been performed under stress induction, which activates PARP1 and thus potentially masks the underlying activities of the other PARP enzymes^3,4,5,6,7^.

Recently, it has been demonstrated that 6-alkyne adenosine (6YnAd), a compound that was previously demonstrated to be suitable for labelling poly(A) tails of mRNAs in mammalian cells^9^, enables sensitive fluorescence profiling of ADP-ribosylated proteins in live cells^10^. In this work, we analysed the labelling efficiency of 6YnAd for the first time using Tandem Mass Tag (TMT)-based quantitative proteomics and found that 6YnAd alone limits substrate coverage, but that inclusion of a similar adenosine analogue, 2-alkyne adenosine (2YnAd), allows a more comprehensive assessment of ADP-ribosylated proteins. We report an integrated chemical proteomics approach that enables the quantitative profiling of ADP-ribosylation of several thousands of endogenous proteins in a robust, reproducible and unbiased manner.

## Results

### 2-alkyne adenosine (2YnAd) treatment results in labelling of ADP-ribosylated proteins

We treated MDA-MB-231 breast cancer cells with equal concentrations of 2YnAd and 6YnAd in parallel. The cells were then lysed and the whole-cell proteome was clicked with a trifunctional capture reagent Azido-TAMRA-Biotin (Suppl. Fig. 1) and resolved on SDS-PAGE. In-gel fluorescence scans revealed qualitatively similar protein labelling profiles for both 2YnAd and 6YnAd treatments (Fig. 1a,b). Western blot analysis of the labelling reactions using anti-pan-ADP-ribose antibody^11^ following affinity enrichment of the labelled proteins on NeutrAvidin-Agarose resin confirmed significant enrichment of ADP-ribosylated proteins in both 2YnAd and 6YnAd live cell incorporations relative to the input control (Fig. 1c,d).

**Figure 1.**
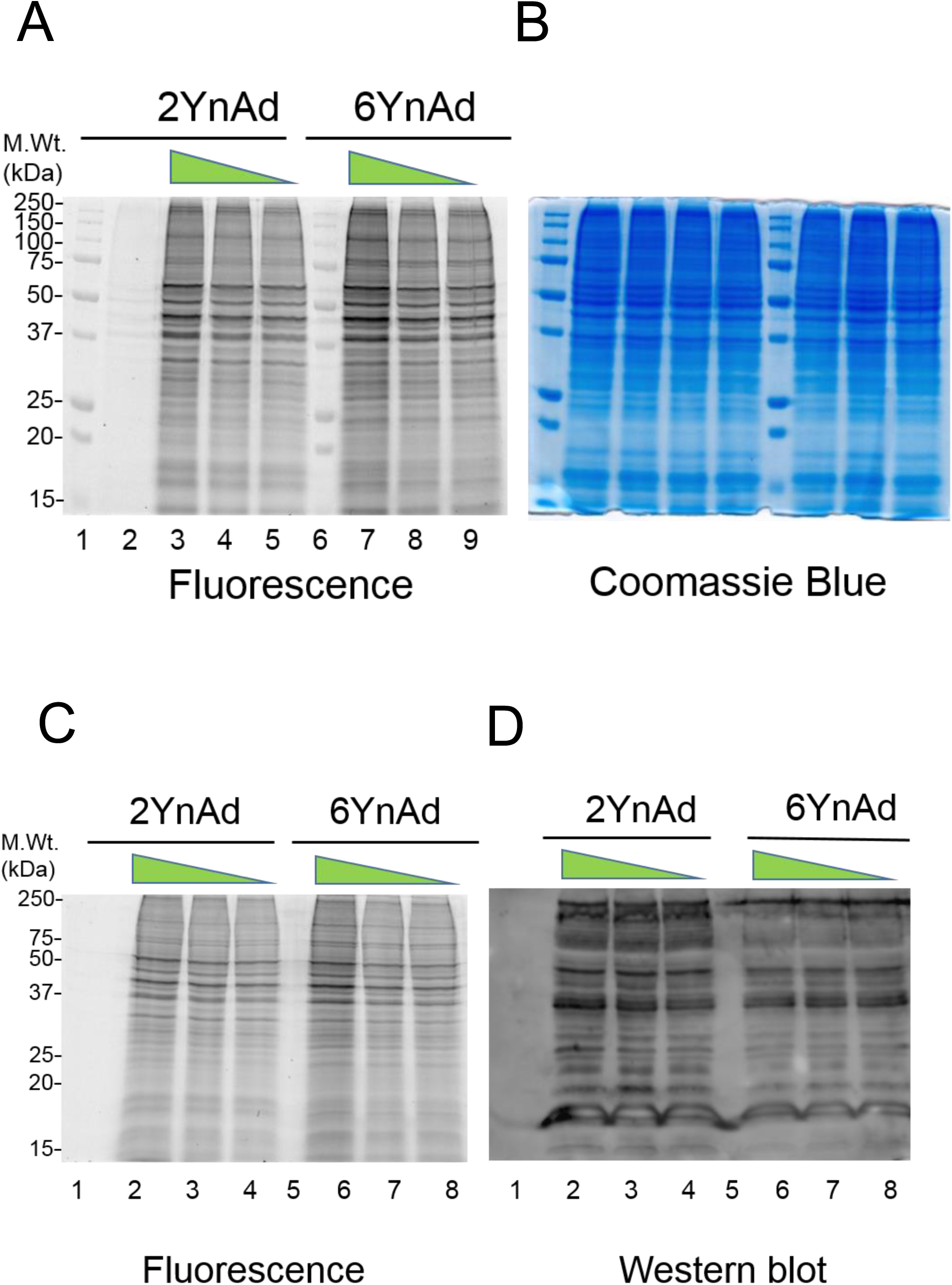
(A) Qualitative assessment of 2YnAd and 6YnAd labelling in MDA-MB-231 cell line by in-gel fluorescence scanning. Lanes 1 and 6: molecular weight marker; lane 2: DMSO control; lanes 3, 4 and 5 (1 mM, 0.5 mM and 0.25 mM 2YnAd respectively) and lanes 7, 8 and 9 (1 mM, 0.5 mM and 0.25 mM 6YnAd respectively). (B) Coomassie blue staining of the same gel. (C) In-gel fluorescence scan following the metabolic incorporation, cell lysis, click chemistry and affinity enrichment. Lane 1: DMSO control; Lanes 2, 3 and 4: 1 mM, 0.5 mM and 0.25 mM respectively of 2YnAd was used for the metabolic labelling; lane 5: empty and lanes 6, 7 and 8: 1 mM, 0.5 mM and 0.25 mM respectively of 6YnAd was used for the metabolic labelling. (D) Western blot of the same gel after electrotransfer of proteins onto a nitrocellulose membrane probed using anti-pan-ADP-ribose antibody (MABE1016, EMD Millipore).

### 2YnAd provides greater coverage of ADP-ribosylated targets than 6YnAd

After establishing the use of the 2YnAd for live cell labelling of ADP-ribosylated proteins, we performed a proteomic comparison of the labelling between 2YnAd and 6YnAd using the highly sensitive tandem mass tags (TMT)-based isobaric tagging for quantitative mass spectrometry^12^. Using the MDA-MB-231 breast cancer cell line as a model system, we first developed a workflow (Fig. 2) that involves the following sequential steps: (i) parallel metabolic incorporation of 2YnAd and 6YnAd, (ii) bioorthogonal tagging of the ADP-ribosylated proteome using copper (I) catalysed azide-alkyne cycloaddition (CuAAC) click reaction, (iii) affinity enrichment of the labelled proteome using the robust biotin-NeutrAvidin pull-down system, (iv) TMT6plex isobaric labelling and (v) liquid chromatography tandem mass spectrometry (LC-MS/MS) analysis.

**Figure 2.**
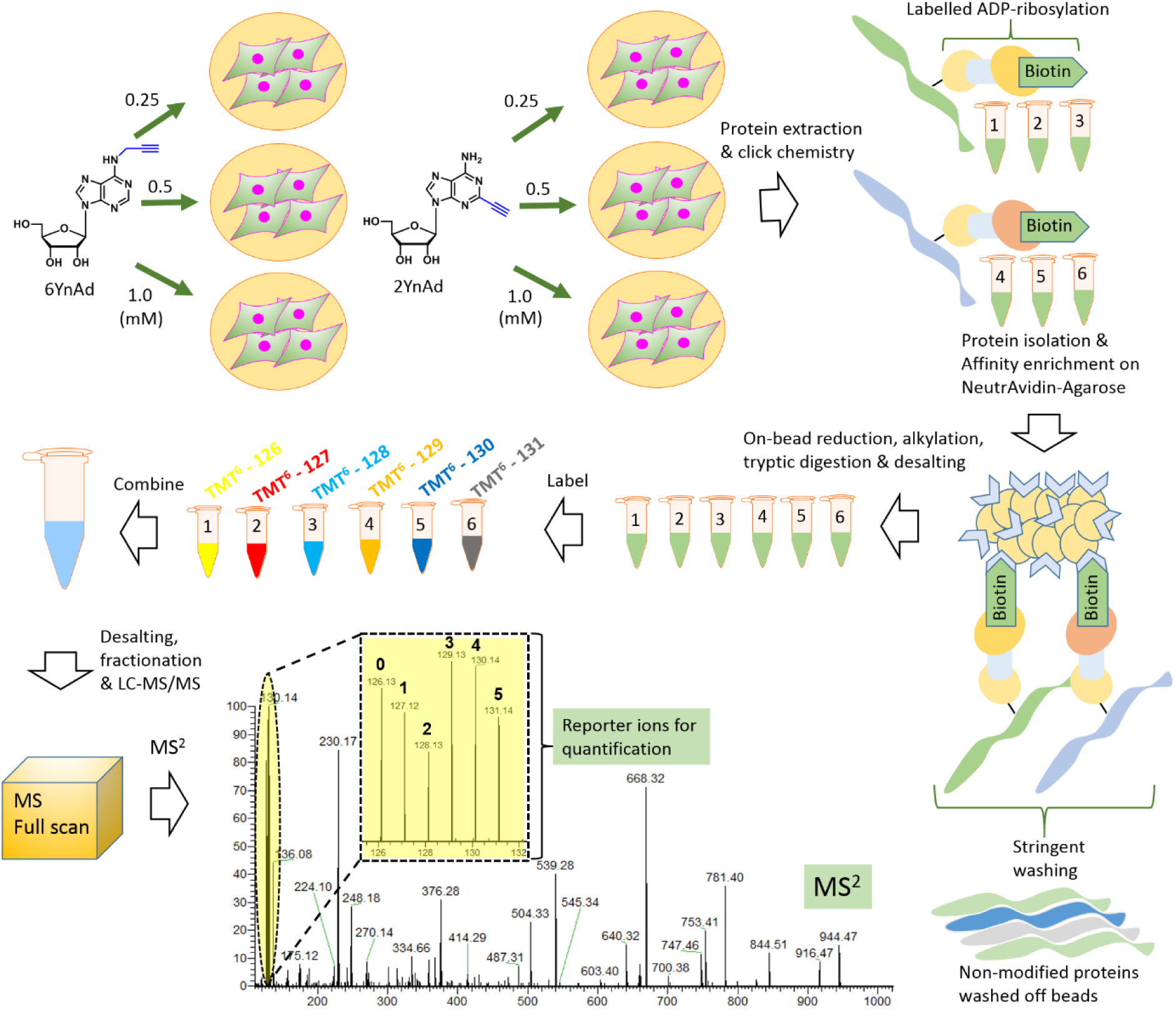
Schematic illustrating the experimental workflow of quantitative comparison of metabolic labelling of 2YnAd and 6YnAd. MDA-MB-231 cells were metabolically labelled with 3 different concentrations of 2YnAd and 6YnAd in parallel. After cell lysis and click reaction with a biotin-containing capture reagent, the labelled proteins are affinity enriched on NeutrAvidin-Agarose resins. Stringent washing at this stage removes non-modified proteins from the beads. After on-bead reduction, alkylation and tryptic digestion, the eluted peptides are desalted and each sample is labelled using a unique TMT6plex channel. After quenching the TMT reactions, the 6 samples are mixed together, fractionated and subjected to nanoflow LC-MS/MS analysis. Quantification of reporter ions from the MS2 spectra provides relative quantification of labelling efficiencies of the two compounds at all three tested concentrations.

Our chemical proteomics workflow for profiling intracellular ADP-ribosylation events identified over 1600 protein targets (Supplementary Dataset File 1, in which proteins previously reported in ADPriboDB^13^ are highlighted in bold), representing a significant improvement in proteomic coverage compared to other published studies despite requiring less protein input (500 µg total protein input before affinity enrichment per condition). Importantly, statistical analysis of the replicate data sets using LIMMA^14^ revealed that specific subsets of protein targets were preferentially labelled with either 2YnAd or 6YnAd treatment (Fig. 3, Suppl. Fig. 2), despite the overall qualitative similarity in the labelling observed in the in-gel fluorescence scans. The 2YnAd was found to be preferentially incorporated, labelling more ADP-ribosylated protein targets than the 6YnAd. Additionally, analysis of the biological pathways of these targets (Suppl. Fig. 3, Supplementary Dataset File 2) specifically revealed a 2YnAd enrichment for proteins involved in proteasome regulation, including several 26S proteasomal subunits.

**Figure 3.**
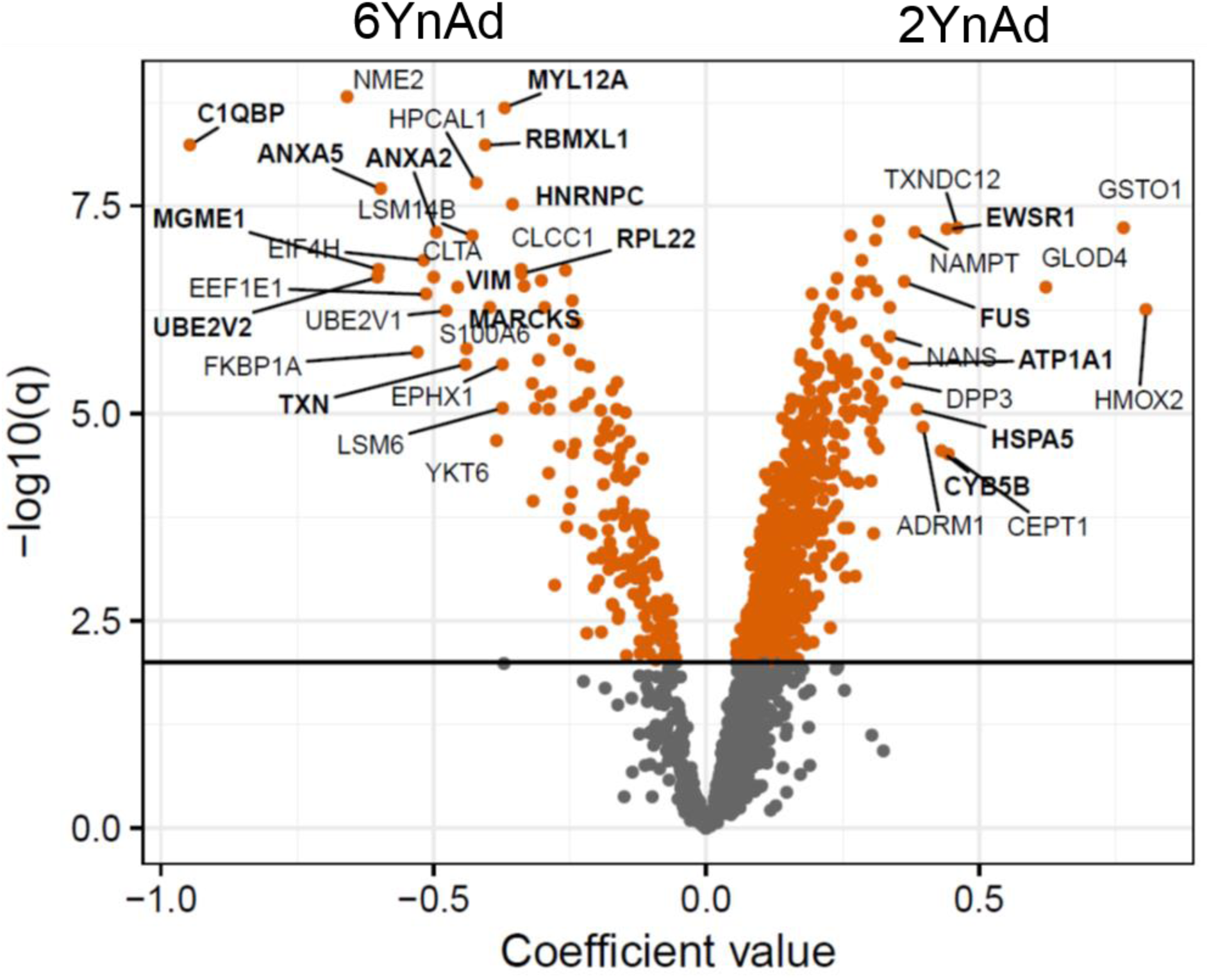
Volcano plot between the cofactor coefficient values (positive values on the x-axis indicate preference for 2YnAd and negative values indicate preference for 6YnAd). Adjusted p values (-log10 scale) following LIMMA analysis are plotted on y-axis with significantly enriched proteins coloured in red. Proteins previously reported in ADPriboDB^22^ are highlighted in bold.

### Dual metabolic labelling using both 2YnAd and 6YnAd elucidates the quantitative response of thousands of ADP-ribosylated proteins to clinical PARP inhibitors

We rationalised that the observed quantitative differences in protein targets labelled by 2YnAd and 6YnAd could be exploited for more comprehensive profiling of the cellular ADP-ribosylome by simultaneously treating cells with both compounds and combining this dual metabolic labelling approach with our quantitative proteomics workflow. Additionally, as several PARP inhibitors have recently emerged as promising cancer therapeutics, we decided to evaluate the dual metabolic labelling strategy for comprehensively and quantitatively profiling global changes to the cellular ADP-ribosylome upon PARP inhibitor treatment (Fig. 4). We treated MDA-MB-231 cells with equimolar concentrations of 2YnAd and 6YnAd to investigate differential labelling of protein ADP-ribosylation for five different concentrations (0, 0.2, 1, 5, 25 µM) of the clinical PARP inhibitors Olaparib and Rucaparib^15^. The samples were combined in a TMT10plex experiment and over 2300 statistically significant hits were identified following LIMMA analysis (Supplementary Dataset File 3), which is the largest experimental candidate data set representing the cellular ADP-ribosylome to date (Fig. 5c). Statistical analysis of the data sets revealed a correlative response in the levels of ADP-ribosylation with respect to Olaparib and Rucaparib treatment, which indicates a congruent set of PARP substrates (Fig. 5a), highlighting the robustness and reproducibility of this integrated chemical biology approach. Several mRNA splicing factors were observed (highlighted in grey rectangle in Fig. 5a) among the most significantly affected proteins to the PARP inhibitor treatments (Fig. 5a, Fig. 5b). Similarly, pathway analysis of the protein targets found to decrease in ADP-ribosylation upon treatment with either PARP inhibitor also identified a preferential enrichment for factors involved in mRNA splicing (Supplementary Dataset File 4, Suppl. Fig. 4).

**Figure 4.**
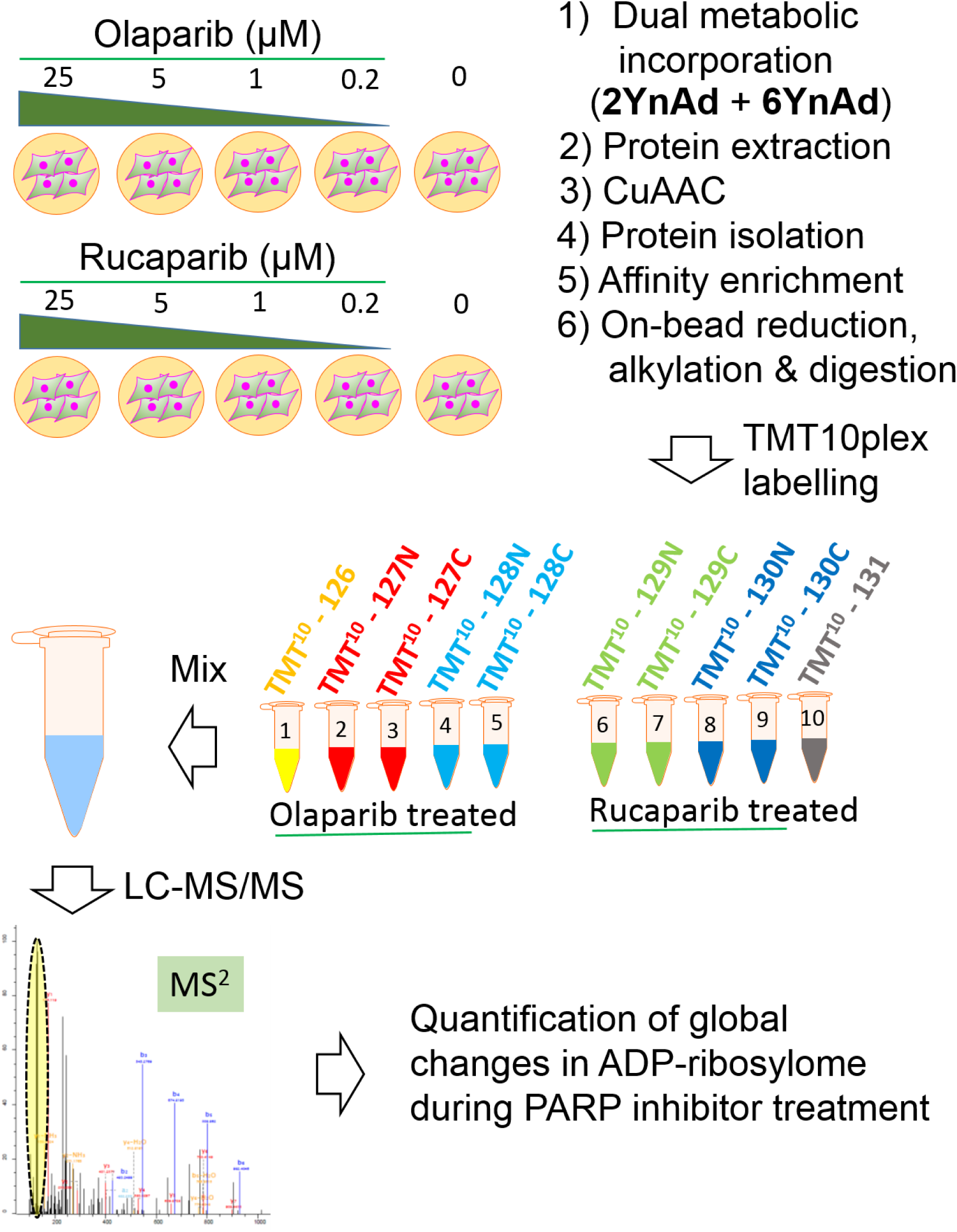
Schematic illustrating the workflow involving parallel treatment of PARP inhibitors Olaparib and Rucaparib followed by dual metabolic labelling of 6YnAd and 2YnAd, click chemistry, affinity enrichment of the labelled proteins on NeutrAvidin-Agarose beads and TMT10plex-based quantification of global changes in the cellular ADP-ribosylome.

**Figure 5.**
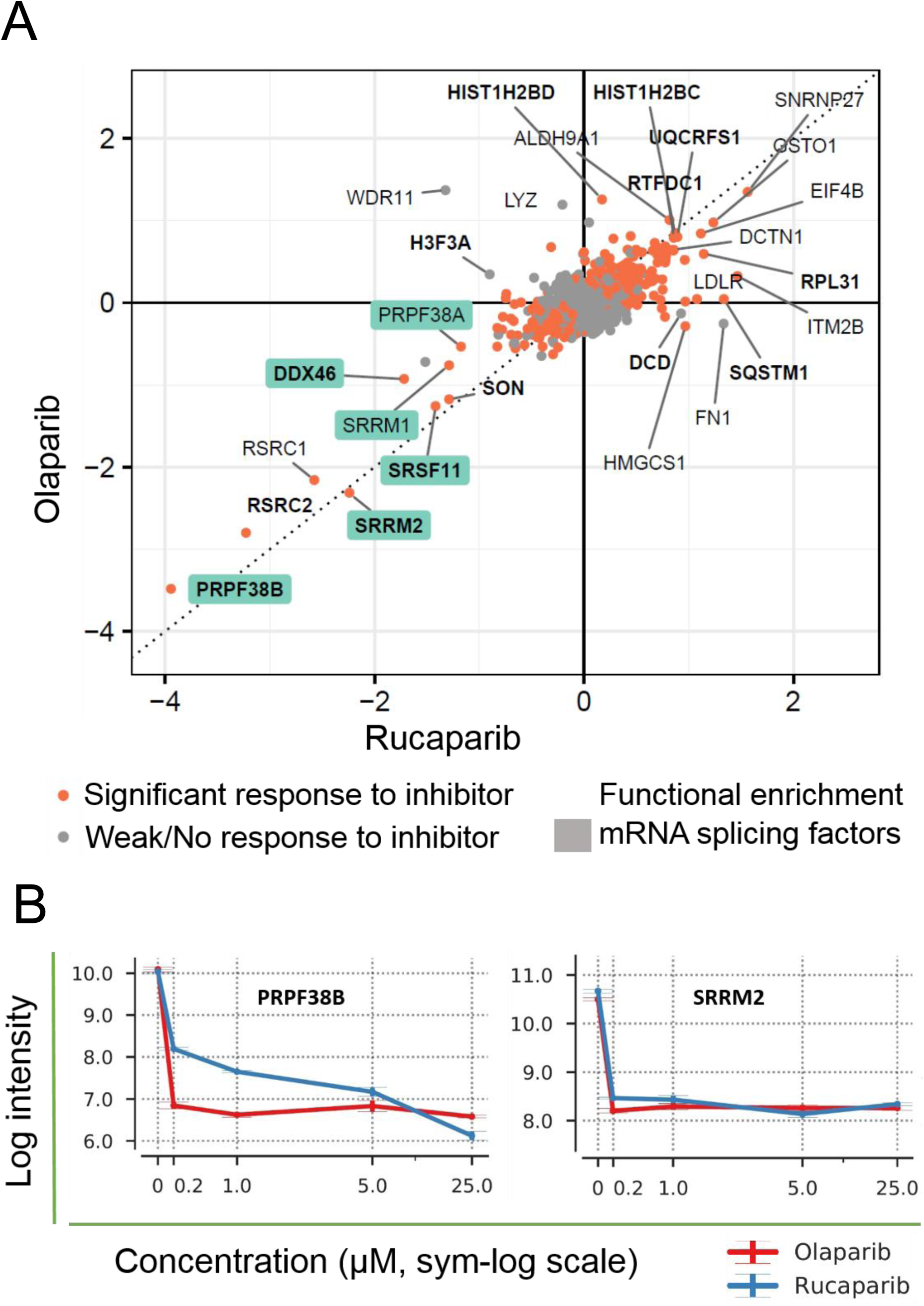
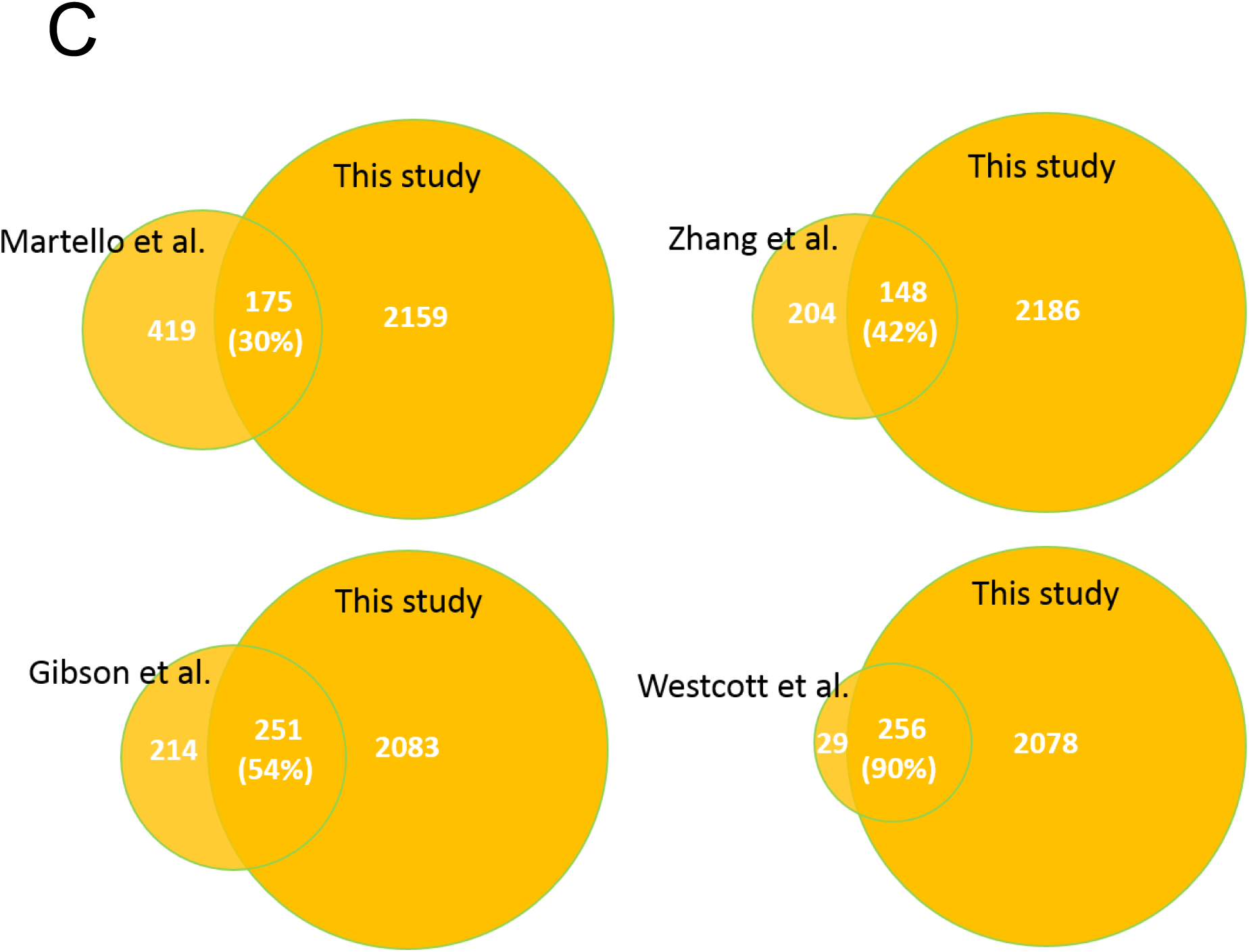
(A) Scatter plot showing mean log2 responses in the level of ADP-ribosylation of endogenous proteins to 25 µM of the inhibitors Olaparib and Rucaparib. Proteins previously reported in ADPriboDB^14^ are highlighted in bold. (B) Quantitative changes in the reporter intensities (log2 scale) of mRNA splicing proteins pre-mRNA splicing factor 38B (PRPF38B) and serine/arginine repetitive matrix protein 2 (SRRM2) as a function of concentration of the inhibitors Olaparib (red) and Rucaparib (blue). (C) Overlap of ADP-ribosylated protein hits between this study and previously published data sets.

## Discussion

Functionalised full-length NAD^+^ analogues have recently emerged as promising tools for fluorescence-based visualisation of ADP-ribosylation in cell-lysates^11,16,17,18^ and in cells^19,20^. Due to the lack of cell-permeability of the full-length NAD^+^ analogues, intracellular labelling was performed by transient permeabilisation of the cell membrane using Triton X-100 surfactant. Although permeabilisation of cell membrane using the surfactant is a powerful approach for imaging applications, successful adaptation of the method to large-scale mass spectrometry (MS) proteomics studies of ADP-ribosylation has not been demonstrated to date. Nevertheless, a chemical genetic discovery method using clickable full-length NAD^+^ analogues has recently enabled selective enrichment and MS profiling of ADP-ribosylated proteins, although the lack of cell permeability of the NAD^+^ analogues has restricted these studies to cell lysates^11^. In a recent advancement in the field, Howard Hang and co-workers reported a chemical biology approach for live cell labelling of ADP-ribosylated proteins using a terminal alkyne functionalised, cell-permeable, NAD^+^-precursor compound, 6-alkyne adenosine (6YnAd)^10^. Although successful labelling of a few hundred ADP-ribosylated proteins was demonstrated using the 6YnAd, previous studies have shown that the corresponding full-length clickable NAD^+^ cofactor (6-alkyne-NAD) is a weaker substrate of PARP1 than endogenous NAD^+16^. This raised the question as to whether the placement of the alkyne group on the NAD^+^ cofactor analogue affects binding and utilisation by the different endogenous PARP enzymes.

We aimed to further develop the metabolic labelling strategy using clickable NAD^+^ precursors, as this approach has the unique advantage of labelling of the ADP-ribosylome directly in the native cellular environment. Since 2-alkyne adenosine (2YnAd), the metabolic precursor for the corresponding full-length clickable NAD^+^ analogue 2-alkyne-NAD^+^, was also reported to efficiently label the poly(A) tails of mRNA transcripts in live cells^21^ similar to the 6YnAd, and recent studies have demonstrated that PARP enzymes are more tolerant to substitutions in this position on NAD^+19,20^, we investigated if 2YnAd can also be used for the metabolic labelling of ADP-ribosylome. Our results (Fig.1) showed for the first time that 2YnAd is also useful for the in-cell labelling of ADP-ribosylated proteins similar to the 6YnAd probe.

As the Western-blot analysis of the labelling by 2YnAd and 6YnAd probes under identical labelling conditions revealed stronger preference for the 2YnAd (Fig. 1d), we decided to apply quantitative proteomics using the highly sensitive TMT isobaric tagging to conduct a head-to-head comparison between of the labelling of protein targets by the two probes. The TMT labelling strategy offers sample multiplexing and importantly the isobaric nature of the tag results in the pooling of peptide signal across all experimental conditions, which increases the sensitivity in detecting lower abundance ADP-ribosylated proteins relative to any non-specific background. Statistical analysis of our TMTsixplex datasets using LIMMA revealed that the 2YnAd was preferentially incorporated, producing more ADP-ribosylated proteins than the 6YnAd probe and suggests the possibility that the endogenous PARP enzymes elicit a differential structural preference for the two NAD^+^ cofactor analogues.

Interestingly, biological pathway analysis of the identified targets revealed enrichment of several proteins involved in proteasome regulation specifically among the 2YnAd targets, including several 26S proteasomal subunits. 26S proteasomal activity has been previously reported to be promoted by ADP-ribosylation^22^. Similarly, a co-operative interaction between the proteasome subunit PSMD14 and BRCA1 has been implicated in cellular responses to DNA double strand breaks^23^, a niche where nuclear PARPs are known to have key functional roles. It has also been suggested that the negatively charged PARylation may serve as a recognition site for the E3 ubiquitin protein ligases, which facilitates subsequent degradation of the protein in the 26S proteasome^24^. However, the precise biological function of ADP-ribosylation in proteasome regulation remains unknown.

Our dual metabolic labelling approach involving simultaneous treatment of cells with equimolar concentrations of 2YnAd and 6YnAd probes and its combination with the TMT10plex quantitative proteomics resulted in the largest candidate experimental dataset of cellular ADP-ribosylome with over 2300 significant proteins. The superior sensitivity of our proteomics technology enabled the systematic profiling of the cellular ADP-ribosylome in an unbiased manner. As an example application, we utilised this methodology to elucidate the first direct and quantitative comparison between the intracellular response in ADP-ribosylation to two clinical PARP inhibitors, Olaparib and Rucaparib. Our findings are in agreement with the recent reports that indicate pivotal functional roles of PARylation in mRNA splicing and modulation of RNAs^25,26,27,28,29^. This finding is further supported by a recent CRISPR screen that identified several high-confidence mRNA splicing factors, including DDX46 and SRSF11 (lower left quadrant in Figure 4A) which resulted in increased sensitivity to the PARP inhibitor Olaparib across several cell lines when mutated^30^.

In summary, we presented a quantitative chemical proteomics approach that combines metabolic labelling using clickable adenosine analogues with TMT isobaric MS to measure ADP-ribosylation in live cells without reduction in PARG activity. We showed that 2YnAd outperforms the previously reported 6YnAd in the metabolic labelling of MDA-MB-231 breast cancer cell line. We demonstrated that our method robustly profiled several thousands of intracellular protein targets, and yielded the largest quantitative proteomics data set measuring the ADP-ribosylome reported to date. The increased sensitivity is the result of the unique combination of the dual metabolic labelling enabled by the clickable NAD^+^ precursors, robust pull-down of modified proteins using NeutrAvidin-Biotin interaction, and the pooling effect resulting from utilising TMT isobaric tagging to multiplex experimental conditions. A head-to-head comparison between two clinical PARP inhibitors revealed a correlative response, which further highlights the sensitivity and robustness of this chemical proteomics platform for studying ADP-ribosylation in live cells. We envision this enabling chemical proteomics technology will transform our ability to systematically characterise the intracellular functions ADP-ribosylation in fundamental and disease biology, including cancer research, bacterial and viral infections and autoimmune diseases. In addition, the approach will be instrumental in determining differences in the targets of established PARP inhibitors and the effects of an emerging range of novel compounds designed to be selective for particular PARP enzymes^31^.

## Methods

### Cell culture

MDA-MB-231 cells (ATCC) were grown in Dulbecco’s Modified Eagle Medium (DMEM) (Gibco) supplemented with 10% foetal bovine serum (FBS) (Gibco) without antibiotics in a humidified incubator with 5% CO_2_ at 37 ^o^C and passaged at every 2-3 days at 80-90% confluency.

### Metabolic labelling and preparation of cell extracts

In a 6-well plate, 4 x 10^5^ cells were treated with 1 ml of fresh media (with 10% FBS) containing 2 µl of appropriate concentration of 6YnAd or 2YnAd (stock solutions in dimethyl sulfoxide [DMSO]) or DMSO control. For labelling experiments in 10 cm plates, 5 ml of fresh media with 10% FBS and 10 µl of the clickable adenosine analogue stock solutions were used. The cells were incubated at 37 ^o^C and 5% CO_2_ for 1 h. The cells were then washed with PBS three times and lysed using 4% SDS lysis buffer (50 mM HEPES pH 7.4, 150 mM NaCl, 4% SDS and 500 U Benzonase) and whole-cell lysates were collected using a cell scraper. Lysates were cleared at 16,000g for 5 min and protein concentrations were estimated using Bio-Rad DC^TM^ protein assay.

### Click chemistry and in-gel fluorescence detection

The cell lysates at 1 mg/ml concentration were treated with freshly premixed click chemistry reaction cocktail (100 µM capture reagent, 1 mM CuSO_4_ solution, 1 mM tris[2-carboxyethyl] phosphine hydrochloride [TCEP], 100 µM [tris(1-benzyl-1H-1,2,3-trazol-4-yl)methyl]amine [TBTA]) for 3 h at room temperature. The following stock solutions of the click chemistry reagents were used. Capture reagent (Azido-TAMRA-Biotin or Azide-PEG3-Biotin (Jena Bioscience)): 10 mM in DMSO; CuSO_4_: 50 mM in MilliQ water; TCEP: 50 mM in MilliQ water; TBTA: 10 mM in DMSO. Proteins were precipitated by adding ice-cold methanol (4 volumes), chloroform (1.5 volumes) and water (3 volumes) and the precipitates were collected by centrifugation at 16,000 g for 5 min. The protein precipitates were then washed twice with ice-cold methanol (10 volumes) and the supernatants were discarded. The protein pellets were air dried for 20 min and suspended in resuspension buffer (4% SDS, 50 mM HEPES pH 7.4, 150 mM NaCl) to 3 mg/ml concentration. Equal volumes of 2X SDS loading buffer (Bio-Rad) containing 10% β-mercaptoethanol were added and the samples were boiled at 95 ^o^C for 10 min and allowed to cool to room temperature. 30 µg of protein was loaded per gel lane (12% SDS Tris-HCl gels) and resolved by SDS-PAGE. The gels were scanned for fluorescence labelling using a GE typhoon 5400 gel scanner.

### Affinity enrichment on NeutrAvidin-Agarose

The dried protein pellets obtained after click chemistry and protein precipitation were suspended in resuspension buffer (4% SDS, 50 mM HEPES pH 7.4, 150 mM NaCl) to 10 mg/ml concentration. In a typical experiment, 500 µg of whole-cell lysate after click chemistry and protein precipitation was resuspended in 50 µl of the resuspension buffer. The samples were subsequently diluted 20 fold with HEPES buffer (50 mM HEPES pH 7.5, 150 mM NaCl) so that the final SDS amount is 0.2%. 50 µl of NeutrAvidin-Agarose beads (Thermo scientific) were washed three times with 0.2% SDS buffer (0.2% SDS 50 mM HEPES pH 7.5, 150 mM NaCl) and added to the samples. The mixtures were rotated on an end-over-end rotating shaker for 1.5 h at room temperature. The beads were then washed 4 times with 1% SDS buffer (1% SDS, 50 mM HEPES pH 7.5, 150 mM NaCl), 4 times with 5 M urea buffer (5 M urea in 50 mM HEPES pH 7.5, 150 mM NaCl) and 4 times with 50 mM freshly prepared ammonium bicarbonate buffer. Each washing was performed with 20 volumes of the washing solutions and centrifugation of the beads between washing steps were carried out at 2,000 g for 1 min. For Western blot analysis, after one round of washing with 1% SDS buffer, the beads were boiled in 4% SDS loading buffer and the proteins were collected by centrifugation. For LC-MS/MS analysis, on-bead reduction, alkylation and tryptic digestion were performed.

### Western blotting

Proteins resolved by SDS-PAGE on 12% Tris-HCl gels were electrotransferred onto a nitrocellulose membrane. Western blotting for ADP-ribosylation was performed using anti-pan-ADP-ribose antibody (MABE1016, lot # 3074935, EMD Millipore) as described earlier with minor modifications.^11^ Briefly, the nitrocellulose membrane after electrotransfer was blocked with freshly prepared 5% BSA in TBS-T for 1 hour at room temperature with constant shaking. The membrane was then incubated with the primary antibody (1:1000 dilution) in 5% BSA in TBS-T for 1 hour at room temperature with constant shaking. The membrane was then washed 4 times for 5 minutes with TBS-T and incubated with Goat-anti-rabbit IgG-HRP secondary antibody (1:5000 dilution) in 5% BSA in TBS-T for 1 hour at room temperature with constant shaking, washed again 4 times for 15 minutes with TBS-T and visualised with Pierce^TM^ ECL Plus Western blotting substrate (Thermo Scientific, Prod # 32132, Lot # SD244941).

### On-bead reduction, alkylation, tryptic digestion and desalting

The beads after affinity enrichment were resuspended in 100 µl of 50 mM ammonium bicarbonate buffer and treated with 3 mM dithiothreitol (DTT) (100 mM stock solution in water) solution for 30 min at room temperature. The beads were washed once with 50 mM ammonium bicarbonate buffer and resuspended in 100 µl of 50 mM ammonium bicarbonate buffer and treated with 10 mM iodoacetamide (IAA) (500 mM stock solution in water) in dark for 30 min at room temperature. The beads were again washed with 50 mM ammonium bicarbonate buffer and resuspended in 100 µl of fresh 50 mM ammonium bicarbonate buffer and treated with 2 µg of sequencing grade modified trypsin (Promega; Product code V5113) at 37 ^o^C overnight. The samples were acidified to pH 3 using formic acid and allowed to stand for 5 min, centrifuged and collected the supernatant. The beads were washed with 0.1% formic acid solution in water, centrifuged and the supernatant were mixed with the previous supernatant. The collected tryptic peptides were desalted on C18 Empore disks on micropipette tip format and the desalted peptide mixture were dried on a speedvac.

### TMTsixplex labelling

MDA-MB-231 cells were fed with three different concentrations (1, 0.5 and 0.25 mM) of 6YnAd and 2YnAd in parallel. After 1 h of incubation, whole-cell lysates were prepared and proteins were quantified. 500 µg of proteins from each condition were subjected to click chemistry using the capture reagent Azido-TAMRA-Biotin. Proteins were precipitated, re-solubilised and subjected to affinity enrichment on NeutrAvidin-Agarose beads. After extensive washings, the beads were subjected to DTT treatment, IAA treatment and overnight trypsinisation. The peptide digests were collected, desalted, dried and re-dissolved in 100 µl of 100 mM triethylammonium bicarbonate (TEAB) buffer (pH 8.5). Each sample was subjected to TMT labelling with a unique TMTsixplex reagent for 1 h at room temperature. After quenching the TMT reactions with hydroxylamine (10 µL of 5% hydroxylamine per reaction for 15 min at room temperature) treatment, the six samples were mixed together, concentrated to complete dryness and re-dissolved in 0.1% formic acid. The sample was desalted on C18 Empore disks on micropipette tip format and the desalted sample was dried on a speedvac.

### Mass-spectrometry analysis of TMT labelled samples

Peptides were reconstituted in 0.1% trifluoroacetic acid and chromatographically resolved using an Ultimate 3000 RSLCnano (Dionex) UHPLC. Peptides were first loaded onto an Acclaim PepMap 100 C18, 3 µm particle size, 100 Å pore size, 20 mm x 75 µm ID (Thermo Scientific) trap column heated to 40 °C, using a loading buffer (2% acetonitrile (MeCN) and 0.05 % trifluoroacetic acid in 97.05 % H_2_O) with a flow rate of 7 µL/minute. Chromatographic separation was achieved using an EASY-Spray column, PepMap C18, 2 µm particles, 100 Å pore size, 500 mm x 75 µm ID (Thermo Scientific) heated to 40 °C. The gradient utilised a flow of 0.3 µL/minute, starting at 98% mobile A (0.1% formic acid, 5% DMSO in H_2_O) and 2% mobile B (0.1% formic acid, 75% acetonitrile, 5% DMSO and 19.9% H_2_O). After 5.5 minutes mobile B was increased to 8% over 0.5 minutes, to 30% over 27 minutes, further increased to 40% in 10 minutes, and finally up to 95% in 2 minutes and held for 4 minutes. Mobile B was reduced back to 5% over 1 minute for the rest of the acquisition.

Data was acquired in real time over 60 minutes using an Orbitrap Fusion Lumos Tribrid mass spectrometer (Thermo Scientific). Eluents were ionised using 2.1 kV spray voltage and a capillary temperature of 275 °C. For all samples MS1 spectra were measured with a resolution of 120,000 at 200 *m/z* and a mass range of 375 to 1500 *m/z*, with a 50 ms maximum injection time, and 4e5 AGC target.

High-resolution HCD MS2 spectra were generated in positive mode for precursor masses using a normalised collision energy of 38%, within a 0.7 *m/z* isolation window, using quadrupole isolation, in top speed mode over a 3 second cycle time, at 50,000 resolution in profile mode, 100 ms maximum injection time and 5e4 AGC target, with an instrument determined scan range beginning at 100 *m/z*.

To ensure quality peptide fragmentation, a number of filters were utilised, including; peptide mono isotopic precursor selection, minimum intensity exclusion of 5e4, and charge state inclusion of 2-5 + ions. To prevent repeat sampling a dynamic exclusion with exclusion count 1, exclusion duration of 30 seconds, +/-10 ppm tolerance window, with isotope exclusion and single charge state selection were used.

### Mass-spectrometry data processing

All raw files were analysed using MaxQuant software (version 1.5.3.30) queried against the Uniprot complete human database (October 2017). The following search parameters were used: Reporter ion MS2 with multiplicity 6plex TMT for the TMTsixplex experiments and 10plex TMT for the TMT10plex experiments, trypsin digestion with maximum 2 missed cleavages, oxidation of methionine and acetylation of protein N-termini as variable modifications, carbamidomethylation of cysteine as a fixed modification, minimum peptide length of 6, protein FDR 0.01. Appropriate correction factors for the individual TMT channels for both peptide N-terminal labelling and lysine side-chain labelling as per the TMT kits used (Thermo Scientific) were configured into the integrated Andromeda search prior to the MaxQuant search.

### PARP inhibitor treatment, dual metabolic incorporation and TMT10plex labelling

MDA-MB-231 cells grown to 80-90% confluency in 10 cm plates were treated with 25, 5, 1, 0.2 and 0 (DMSO control) µM concentrations of the PARP inhibitors Olaparib (10 mM stock solution in DMSO) and Rucaparib (10 mM stock solution in DMSO) in parallel. All treatments were performed in 10 ml DMEM with 10% FBS and the cells were placed in an incubator with 5% CO_2_ and 37 ^o^C for 1 h. After 1 h, the media were removed and the cells were supplemented with fresh complete media (5 ml) containing 0.25 mM concentrations of both 6YnAd (500 mM stock solution in DMSO) and 2YnAd (500 mM stock solution in DMSO) continued incubation for one more hour. The cells were washed with PBS three times and whole-cell lysates were prepared and proteins were quantified. 500 µg protein per each condition were subjected to click reactions using Azide-PEG3-Biotin capture reagent. Proteins were precipitated using the methanol/chloroform/water system and air-dried precipitates were resuspended in the pull-down buffer (50 mM HEPES, pH 7.4, 0.2% SDS, 150 mM NaCl) and subjected to affinity enrichment on NeutrAvidin-Agarose beads. After extensive washings, the beads were subjected to DTT treatment, IAA treatment and overnight trypsinisation. The peptide digests from the 10 conditions were collected, desalted, dried and re-dissolved in 100 µl of 100 mM triethylammonium bicarbonate (TEAB) buffer (pH 8.5). Each sample was subjected to TMT labelling with a unique TMT10plex reagent for 1 h at room temperature. After quenching the TMT reactions with hydroxylamine treatment, the 10 samples were mixed together, concentrated to complete dryness and re-dissolved in 0.1% formic acid solution in water. The sample was desalted on C18 Empore disks on micropipette tip format and the desalted sample was dried on a speedvac. The sample was dissolved in 300 µl of 0.1% TFA solution in water and subjected to fractionation using High pH Reverse-Phase Peptide Fractionation Kit (Thermo scientific) to 8 fractions and dried on a speedvac.

### Pre-processing of MaxQuant output for data analysis

Columns corresponding to the corrected reporter intensity were extracted from the proteinGroups.txt files in MaxQuant output. Potential contaminants, proteins only identified by site, reverse proteins and proteins containing less than two peptides were discarded. Out of the remaining proteins, only proteins that have non-zero intensities across all reporters and replicates were kept (1604 proteins in 2YnAd-6YnAd dataset, 2334 in the inhibitor dataset). Intensities of the remaining proteins were normalised using TMM method^32^ and converted to log2-per-million values. The resulting data was then compared with previously reported annotations of ADP-ribosylated proteins available in ADPriboDB (v1.0)^13^, which was augmented to include annotations reported since 2015 (v2.0; unpublished), after prefiltering the database to contain only annotations derived from human datasets.

### LIMMA linear model design

Both cofactor preference and inhibitor treatment datasets were analysed using LIMMA method. The linear methods for each of these datasets were designed separately and described below.

#### 2YnAD/6YnAD

The normalised intensities are assumed to be modelled by the following effects in a format described below:

> log2(normalised intensity) ~ [Cofactor effect, Sum encoding] + [log(concentration) effect] + [Replicate effect] + Intercept.

We used the following encoding for the effects.

1. **Cofactor effect** (i.e. the main measured effect). Encoded by +1 in reporters corresponding to 2YnAd and −1 in reporters corresponding to 6YnAd (e.g. sum-to-one encoding). Positive coefficients to this factor therefore indicate 2YnAd preference, and negative values indicate 6YnAd preference.
2. **log(concentration) effect**: encoded as log values of the concentrations in mM. Effect is assumed to be the same for both cofactors. Higher values indicate stronger response.
3. **Replicate**: 1-of-K encoding (minus one redundant term absorbed in intercept). This term accounts for technical-replicate associated bias in protein intensities.

The complete design matrix is available as Supplementary Table 1.

#### PARP Inhibitor dataset

For this dataset we have applied the linear modelling only on the data points corresponding to 25µM of inhibitor and 0 µM (reporters 0, 4, 5 and 9) only, not taking the data for intermediate concentrations into account. The following linear model was assumed.

Log(intensity) ~ [Inhibitor preference effect] + [Inhibition effect] + [Replicate bias] + [Intercept].

The three factors have been encoded as follows:

1. **Inhibitor preference effect**. One of the two major effects in this experiment. This factor is designed to capture the effect of a particular inhibitor on the intensity of the protein. We encoded this using the following map: −1 = Olaparib, +1 = Rucaparib and 0 = No inhibitor (i.e. concentration = 0 µM). In this way, positive values correspond to protein having to higher intensity upon Rucaparib treatment and negative values corresponding to higher intensity upon Olaparib treatment.
2. **Inhibition effect**. This term corresponds to the second major effect in the experiment, which captures the general effect of inhibition to the protein that does not depend on the inhibitor. It was encoded as 1 (in cases where inhibitor is present) and 0 otherwise. This way values below than zero correspond to decrease in intensity upon inhibition.
3. Replicate bias: This term accounts for technical-replicate associated bias in protein intensities. This was encoded with one-in-K encoding with one redundant term absorbed to intercept.

The complete design matrix is available in Supplementary Table 2.

### LIMMA processing and statistical testing

We have used a similar procedure to the one described by Kammers et al^33^. The linear models were encoded as described above. LIMMA analysis was run with the trend parameter set to TRUE (commonly referred to as LIMMA-trend method). After the model has been fit, we have performed statistical testing on the effect coefficients in the model using the equations provided by Kammers et al. Specifically, we have tested the null hypothesis that cofactor effect coefficient is zero in 2YnAd/6YnAd dataset, and the null hypotheses that inhibitor preference effect and inhibition effect coefficients are not zero (separately) in inhibitor treatment dataset. The p values were adjusted with Benjamini-Hochberg method. Any adjusted p values below 0.01 were deemed significant and annotated in the plots and output. In order to generate debiased plots, we have subtracted the values for bias coefficients (e.g. replicate bias and intercept) from the normalised log2 intensities.

### Pathway enrichment analysis

Biological pathway enrichment analysis was performed against Reactome database^34^ using the R packages ReactomePA^35^ and clusterProfiler^36^ using the default settings, showing the most descriptive categories with the “by=rowPercentage” parameter set.

## Data Availability

The tandem mass spectrometry datasets generated during the current study are available via the PRIDE database (project accession: PXD011631). All of the data analysed during this study are included in this published article (and its Supplementary Information files).

## Acknowledgements

This work was supported by GlaxoSmithKline through the Imperial College London Engineering Medicines Laboratory Project (K.A.K, D.O.H and P.A.D) and the UK Biotechnology and Biological Sciences Research Council Doctoral Training Partnership award (S.L). A.P.S. and A.B. are supported by the Francis Crick Institute, which receives its core funding from Cancer Research UK (FC001999), the UK Medical Research Council (FC001999) and the Wellcome Trust (FC001999).

## Author contribution

K.K and P.A.D conceived of the study; K.K performed experiments; S.L and P.A.D performed statistical analysis of data; K.K, S.L, A.J.B, A.P.S, D.O.H, A.K.L.L and P.A.D contributed to experimental design, data analysis and data interpretation. K.K and P.A.D wrote the manuscript with input from all authors.

