## Supplemental Information for "An Integrated Chemical Proteomics Approach for Quantitative Profiling of Intracellular ADP-Ribosylation"

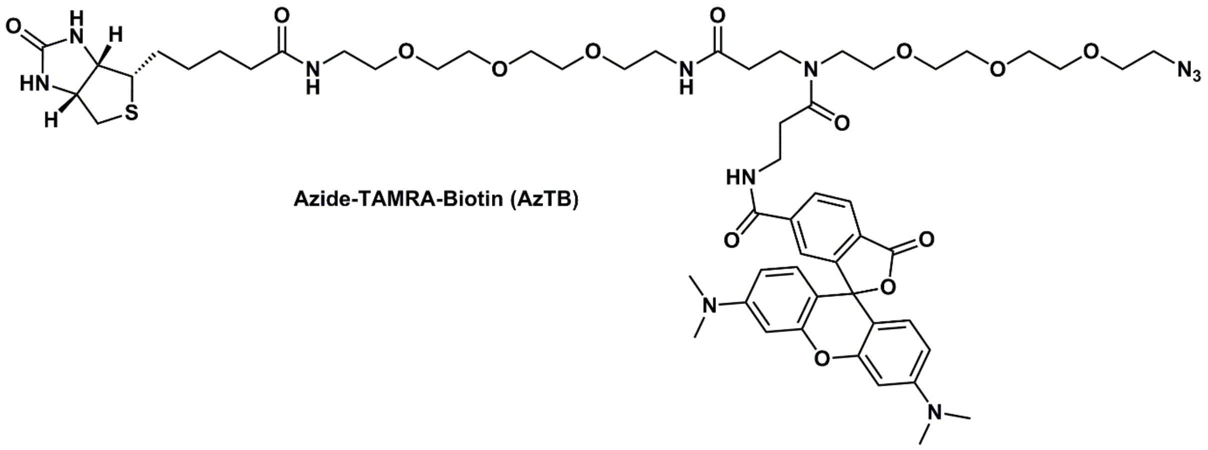

**Azide-TAMRA-Biotin (AzTB)**

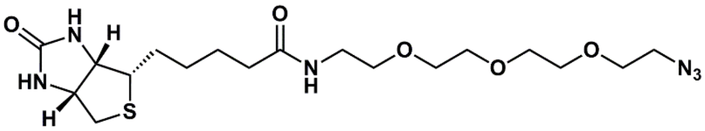

**Azide-PEG3-Biotin**

***Suppl. Fig. 1*.** Chemical structures of the capture reagents used in this study.

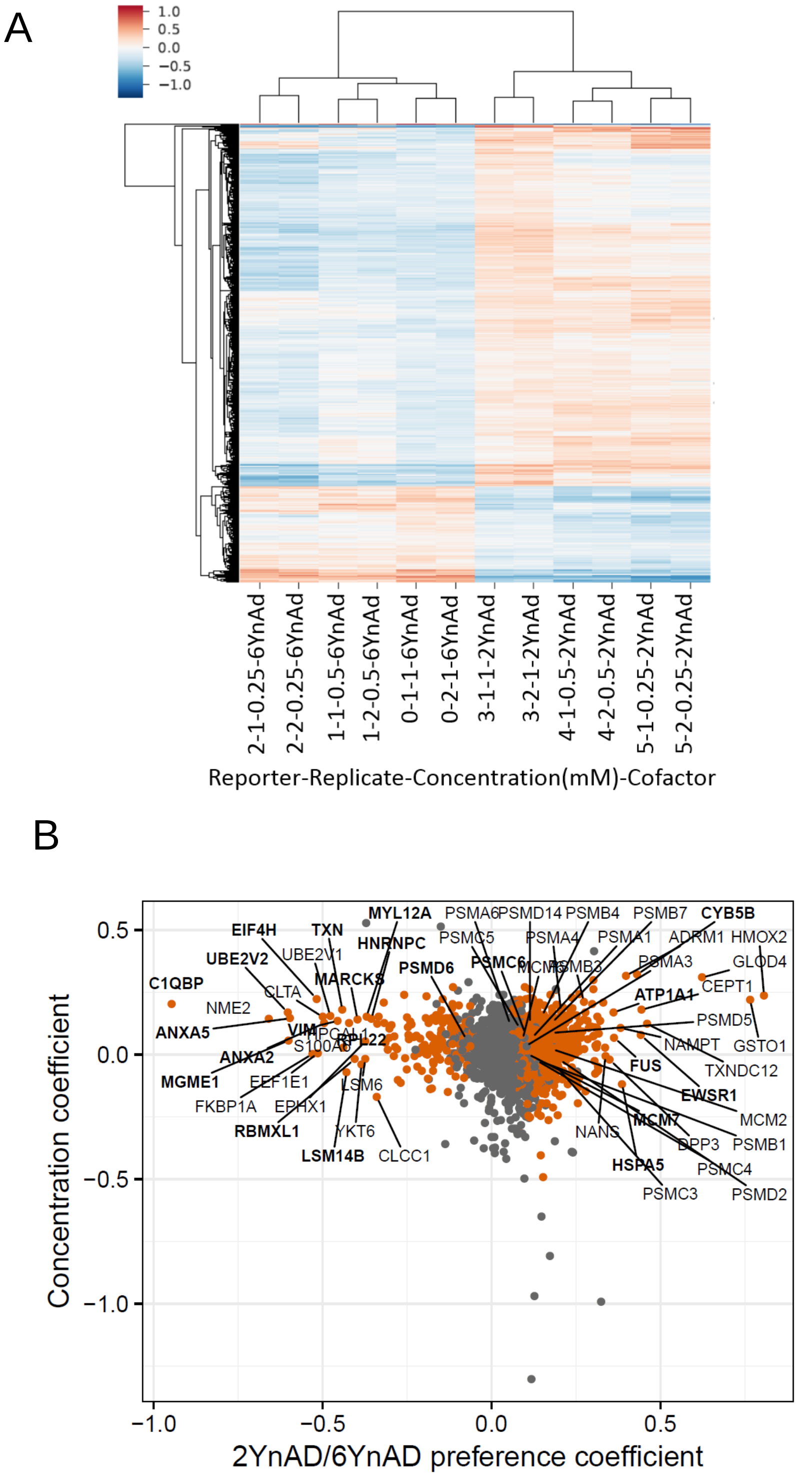


***Suppl. Fig. 2.*** (A) Heatmap indicating mass spectrometry intensities across all three concentrations after subtracting bias terms (e.g. replicate bias and intercept). Red colour indicates higher intensity compared to mean across row. (B) Scatter plot of the value relationship with concentration effect coefficient of the TMTsixplex labelling experiment following LIMMA statistical analysis. Entries in the updated (unpublished) version of the ADPriboDB are highlighted in bold.

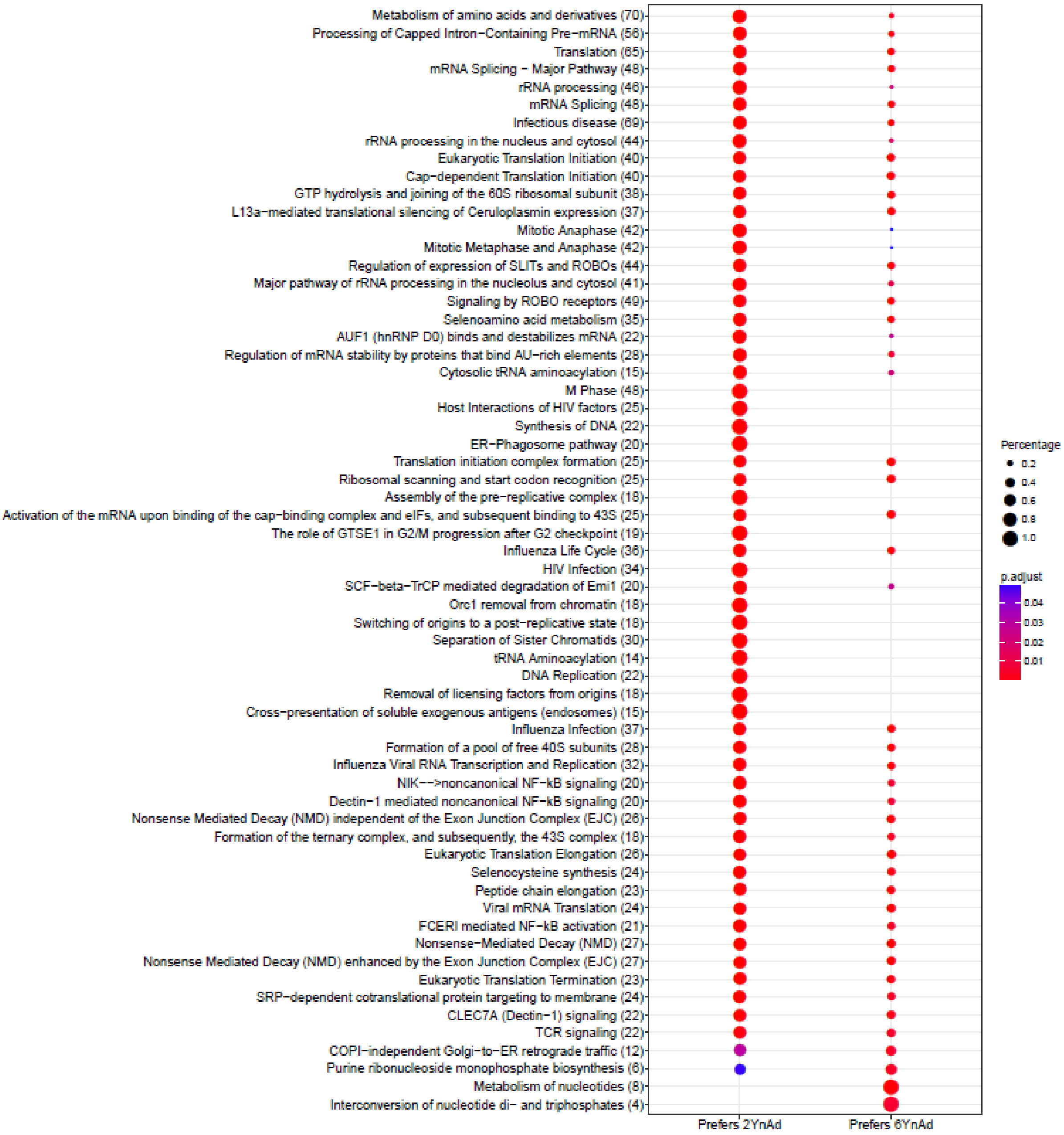


***Suppl. Fig. 3***. Biological pathway enrichment analysis dot-plots of the preferred targets of 2YnAd and 6YnAd.

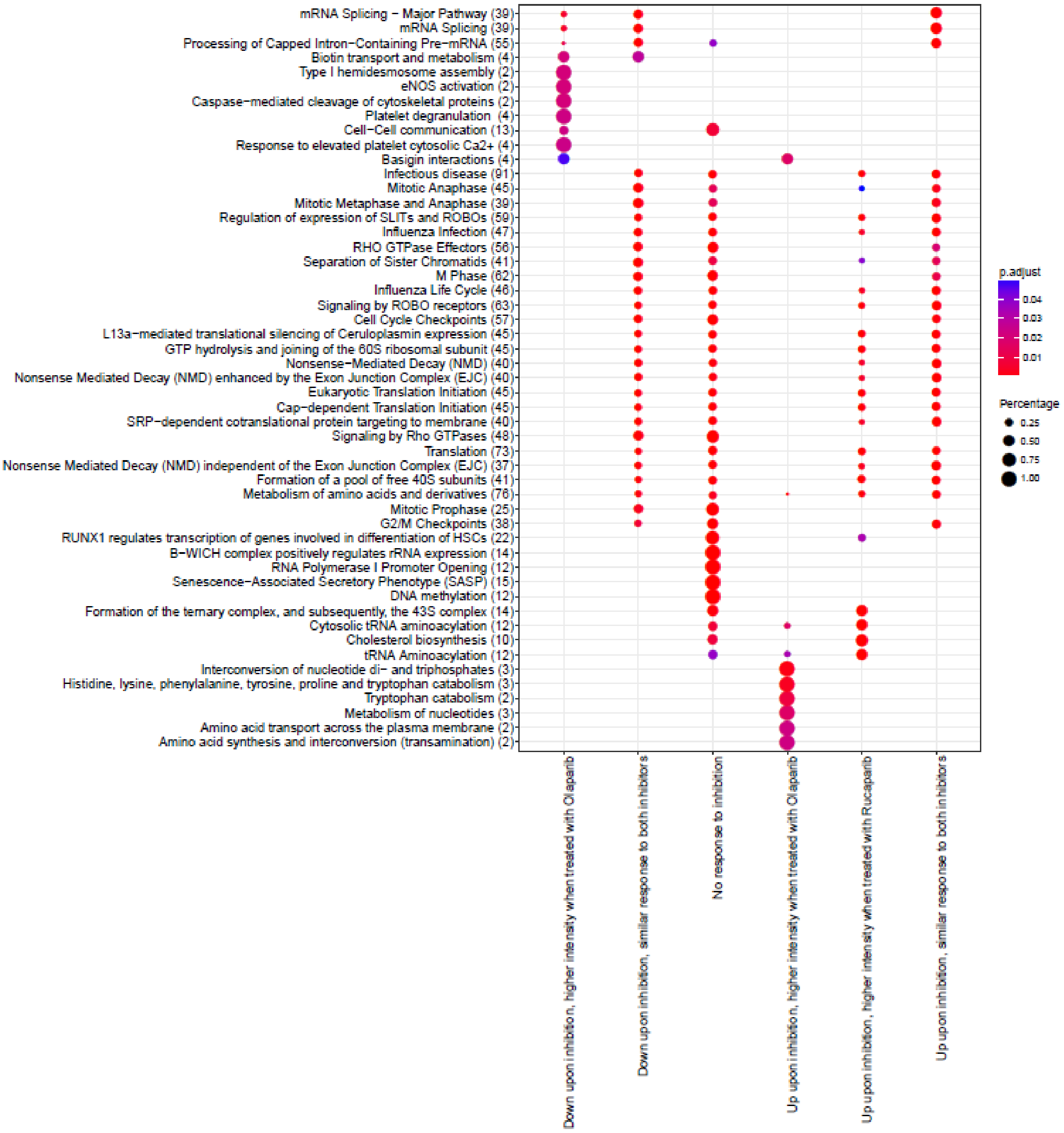


***Suppl. Fig. 4***. Biological pathway enrichment analysis dot plots of the TMT10plex labelling experiment.

(A)

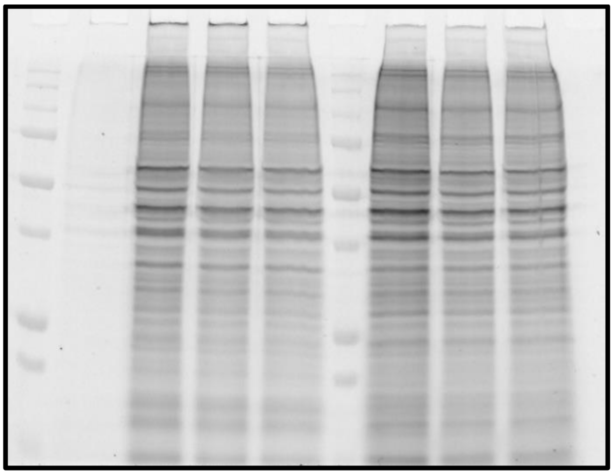

(B)

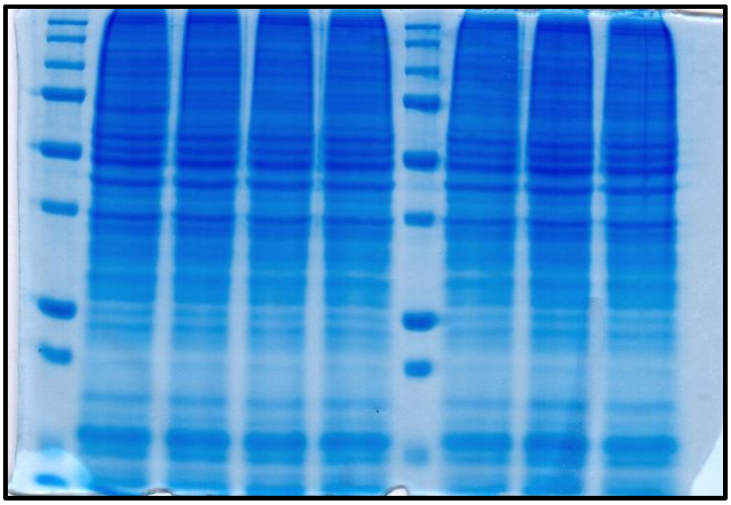

**(C)**

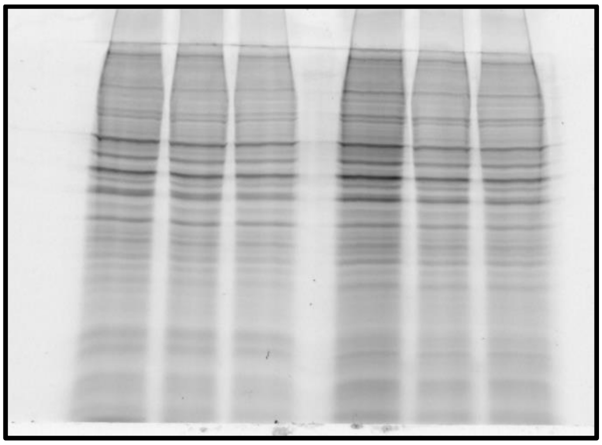

**(D)**

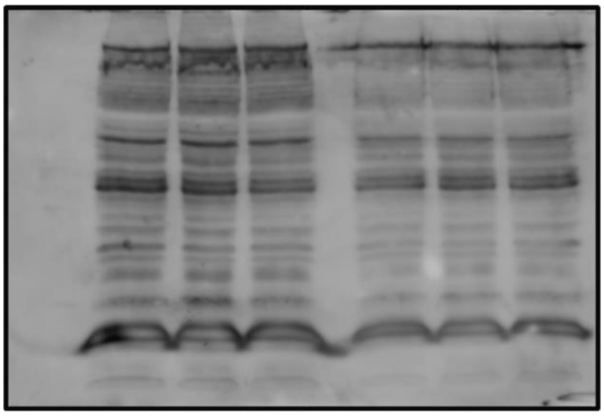

***Suppl. Fig. 5***. Full-length gels and blot used in Figure 1.

### Supplementary Table 1

Design matrix for 2YnAD/6YnAD dataset

| Headers | | | | Design matrix | | | |
|---|---|---|---|---|---|---|---|
| **Reporter** | **Replicate** | **Concentration _mM** | **Cofactor** | **Intercept** | **C(Cofactor, Sum)[S.2YnAd]** | **C(Replicate)[T.2]** | **np.log(Concentration_mM)** |
| **0** | **1** | **1** | **6YnAd** | 1 | -1 | 0 | 0 |
| | **2** | **1** | **6YnAd** | 1 | -1 | 1 | 0 |
| **1** | **1** | **0.5** | **6YnAd** | 1 | -1 | 0 | -0.693147 |
| | **2** | **0.5** | **6YnAd** | 1 | -1 | 1 | -0.693147 |
| **2** | **1** | **0.25** | **6YnAd** | 1 | -1 | 0 | -1.386294 |
| | **2** | **0.25** | **6YnAd** | 1 | -1 | 1 | -1.386294 |
| **3** | **1** | **1** | **2YnAd** | 1 | 1 | 0 | 0 |
| | **2** | **1** | **2YnAd** | 1 | 1 | 1 | 0 |
| **4** | **1** | **0.5** | **2YnAd** | 1 | 1 | 0 | -0.693147 |
| | **2** | **0.5** | **2YnAd** | 1 | 1 | 1 | -0.693147 |
| **5** | **1** | **0.25** | **2YnAd** | 1 | 1 | 0 | -1.386294 |
| | **2** | **0.25** | **2YnAd** | 1 | 1 | 1 | -1.386294 |

### Supplementary Table 2

Design matrix for inhibitor dataset.

**Headers** **Design matrix**

| Reporter | Replicate | Concentration_uM | Inhibitor | Inhibitor_binary | Inhibitor_added | C(Inhibitor_binary, contrast)[custom0] | C(Inhibitor_added)[T.True] | C(Replicate)[T.2] | C(Replicate)[T.3] | Intercept |
|---|---|---|---|---|---|---|---|---|---|---|
| **0** | **1** | 25 | Olaparib | Olaparib | TRUE | -1 | 1 | 0 | 0 | 1 |
| | **2** | 25 | Olaparib | Olaparib | TRUE | -1 | 1 | 1 | 0 | 1 |
| | **3** | 25 | Olaparib | Olaparib | TRUE | -1 | 1 | 0 | 1 | 1 |
| **4** | **1** | 0 | Olaparib | NA | FALSE | 0 | 0 | 0 | 0 | 1 |
| | **2** | 0 | Olaparib | NA | FALSE | 0 | 0 | 1 | 0 | 1 |
| | **3** | 0 | Olaparib | NA | FALSE | 0 | 0 | 0 | 1 | 1 |
| **5** | **1** | 25 | Rucaparib | Rucaparib | TRUE | 1 | 1 | 0 | 0 | 1 |
| | **2** | 25 | Rucaparib | Rucaparib | TRUE | 1 | 1 | 1 | 0 | 1 |
| | **3** | 25 | Rucaparib | Rucaparib | TRUE | 1 | 1 | 0 | 1 | 1 |
| **9** | **1** | 0 | Rucaparib | NA | FALSE | 0 | 0 | 0 | 0 | 1 |
| | **2** | 0 | Rucaparib | NA | FALSE | 0 | 0 | 1 | 0 | 1 |
| | **3** | 0 | Rucaparib | NA | FALSE | 0 | 0 | 0 | 1 | 1 |

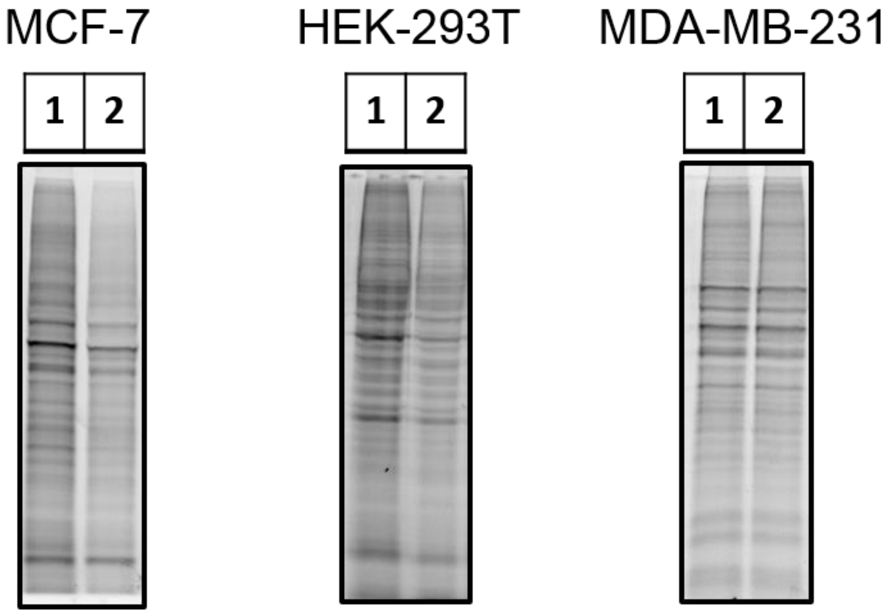


***Suppl. Fig. 6***. 2YnAd labelling in MCF-7, HEK-293T and MDA-MB-231 cell lines. Lane 1 (0.5 mM) and lane 2 (0.25 mM) probe.

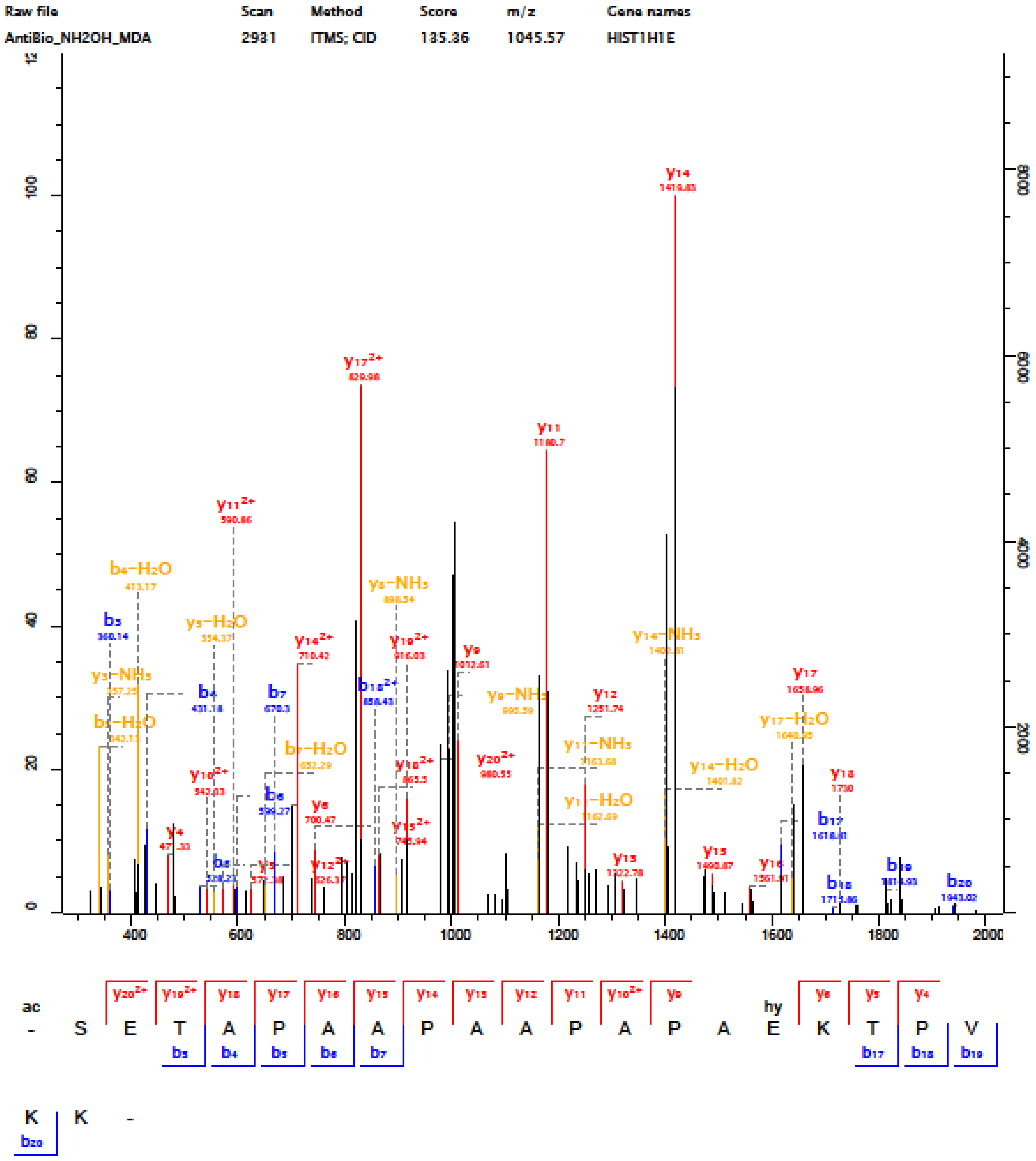

Raw file
AntiBio_NH2OH_MDA
Scan
2931
Method
ITMS; CID
Score
135.36
m/z
1045.57
Gene names
HIST1H1E
0
20
40
60
80
100
2000
4000
6000
8000
400
600
800
1000
1200
1400
1600
1800
2000
ac
hy
- S E T A P A A P A A P A P A E K T P V
K K -

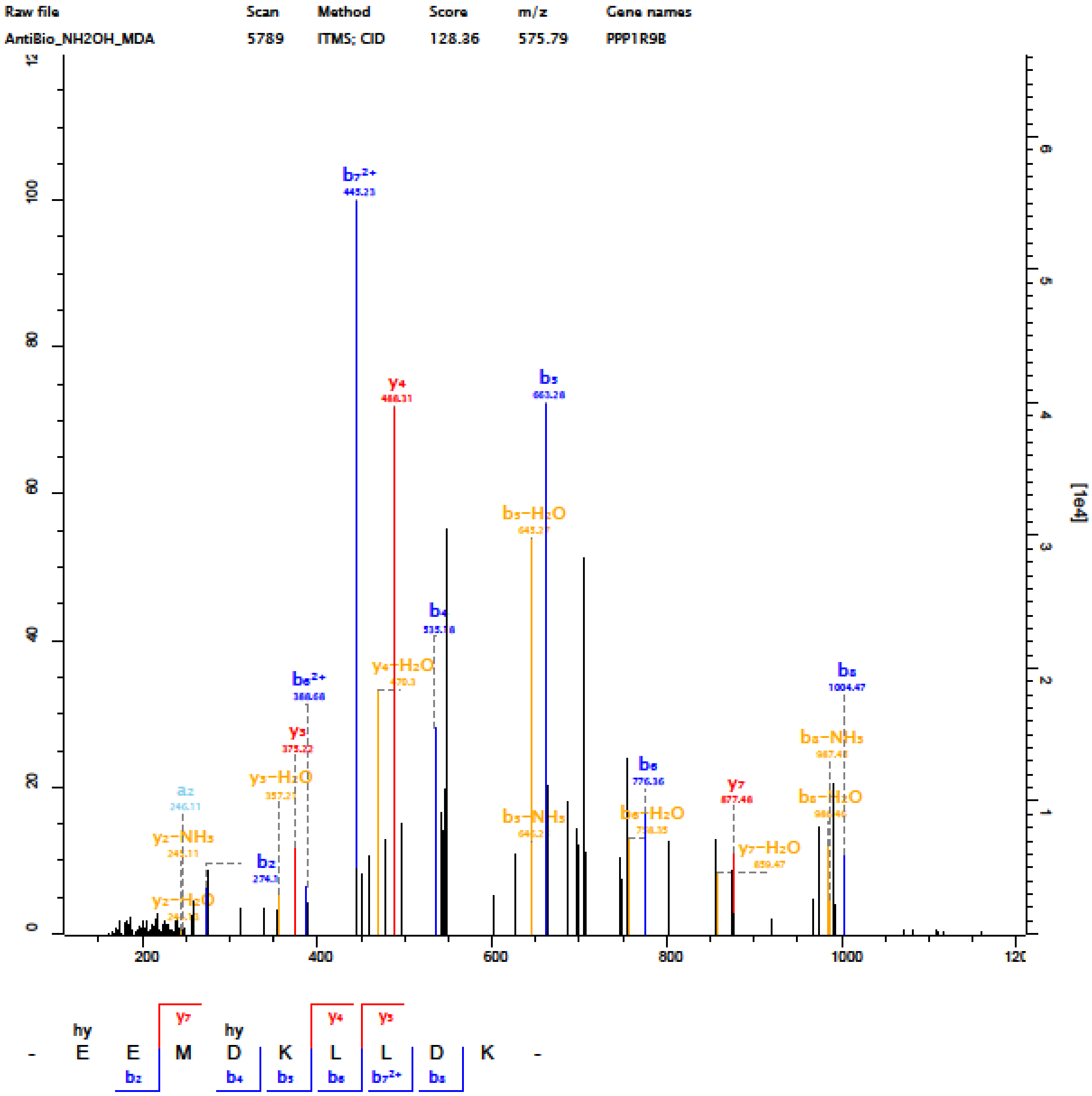
Raw file
AntiBio_NH2OH_MDA
Scan
5789
Method
ITMS; CID
Score
128.36
m/z
575.79
Gene names
PPP1R9B
[1e4]
b7²⁺
445.23
y4
488.31
b5
663.28
b5−H2O
645.27
b4
535.18
y4−H2O
b8
1004.47
b6²⁺
y3
375.22
y3−H2O
a2
y2−NH3
b2
y2−H2O
b6
776.36
y7
877.48
y7−H2O
b8−NH3
b8−H2O
b5−NH3
b6−H2O
- E E M D K L L D K -
hy
y7
y4
y3
b2
b4
b5
b6
b7²⁺
b8

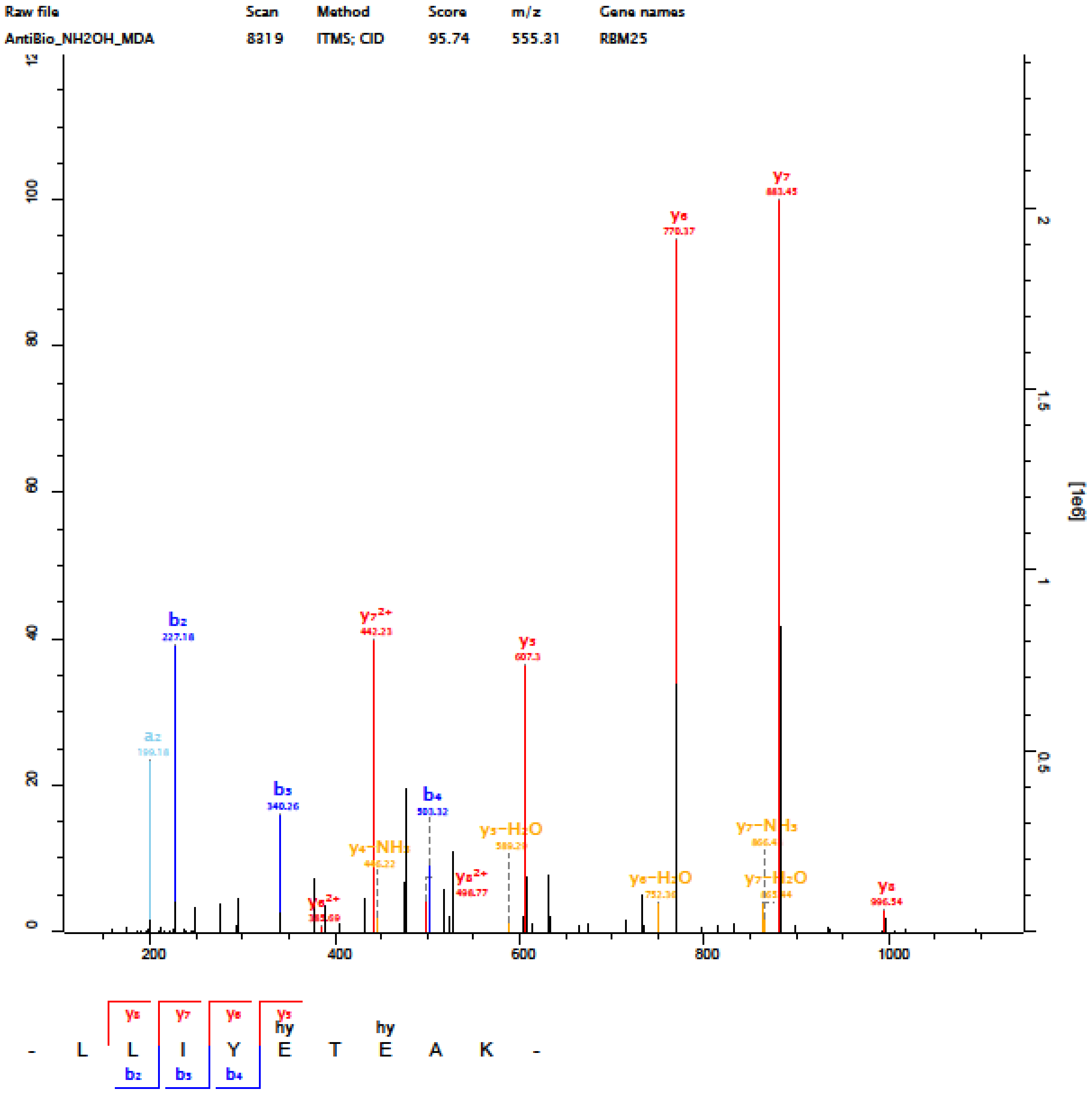


**_Suppl. Fig. 7_**. Representative MS/MS spectra of modified peptides. Affinity enriched ADP-ribosylated peptides were treated with hydroxylamine to hydrolyse the PARylation to size-reduced, MS-compatible, adducts at the modification sites with a characteristic mass signature of +15.0109 Da. The modification sites in the tryptic peptide sequence (below each spectrum) were indicated using the symbol “hy”.
